# Antibiotic Treatment Drives the Diversification of the Human Gut Resistome

**DOI:** 10.1101/537670

**Authors:** Jun Li, Elizabeth A. Rettedal, Eric van der Helm, Mostafa Ellabaan, Gianni Panagiotou, Morten O.A. Sommer

## Abstract

Despite the documented antibiotic-induced disruption of the gut microbiota, the impact of antibiotic intake on strain-level dynamics, evolution of resistance genes, and factors influencing resistance dissemination potential remains poorly understood. To address this gap we analyzed public metagenomic datasets from 24 antibiotic treated subjects and controls, combined with an in-depth prospective functional study with two subjects investigating the bacterial community dynamics based on cultivation-dependent and independent methods. We observed that short-term antibiotic treatment shifted and diversified the resistome composition, increased the average copy number of antibiotic resistance genes, and altered the dominant strain genotypes in an individual-specific manner. More than 30% of the resistance genes underwent strong differentiation at the single nucleotide level during antibiotic treatment. We found that the increased potential for horizontal gene transfer, due to antibiotic administration, was ∼3-fold stronger in the differentiated resistance genes than the non-differentiated ones. This study highlights how antibiotic treatment has individualized impacts on the resistome and strain level composition, and drives the adaptive evolution of the gut microbiota.

## Introduction

The human intestines are densely populated by diverse microbial inhabitants, which collectively constitute the gut microbiota. About 1,000 prevalent bacterial species colonize the human gastrointestinal tract, playing a pivotal role in health and disease of the host. Besides influencing physiology of the digestive tract, the gut microbiota also affects development, immunity, and metabolism of the host [1]. External forces, including antibiotic treatment or dietary intake, shape the composition of the gut microbiota with the potential for rapid changes, thereby affecting the microbe-host homeostasis [2,3].

Antibiotics have been widely used since the Second World War resulting in dramatic benefits to public health [4]. However, the rapid increase of antibiotic resistance (AR) has become an escalating worldwide issue [5]. Antibiotic-resistant pathogens lead to treatment failure and contribute to increasing morbidity, mortality, and healthcare costs. Over 70% of bacteria causing hospital-acquired infections have antibiotic resistance towards at least one common antibiotic for treatment [6]. Previous metagenomic studies have revealed the influence of antibiotic administration on the gut microbiota in various ways, including 1) altering the global taxonomic and functional composition or the diversity of the gut microbiota [7–11] 2) increasing the abundance of bacteria resistant to the administered antibiotic [12] 3) expanding the reservoir of resistance genes (resistome) [13] or 4) increasing the load of particular antibiotic resistance genes (ARGs) [11,14]. The disruptive effects of antibiotic treatment on gut microbiota can be transient or long-lasting [15]. Nevertheless, there is a paucity of information about how the resistome structure shifts and how the genotype of ARGs evolve and differentiate when the microbiome is challenged with antibiotics. In addition, how antibiotic exposure influences strain-level variation within the gut microbiome remains poorly understood.

Antibiotic resistance can be acquired through point mutations (*de novo* evolution) or horizontal gene transfer (HGT) [16]. *De novo* resistance mutations can modify the antibiotic cellular targets or alter the expression of antibiotic resistance genes and therefore alter the resistance levels of the bacterial strain harboring them [17]. Unlike *de novo* resistance mutations, horizontal gene transfer allows bacteria to adapt more rapidly to an environment containing antibiotics [16]. Furthermore, the gastrointestinal tract, densely populated with bacteria, enables the gut microbiota to act as a resistance reservoir which likely contributes to the spread of ARGs to opportunistic pathogenic bacteria [18-20]. The exchange of ARGs has been documented to occur in the human gut between strains carrying vancomycin and sulfonamide resistance genes in *Enterococcus faecium* and *Escherichia coli*, respectively [21,22]. Recently, the *in situ* HGT of ARGs in the infant gut was described [23]. However, factors triggering the HGT of ARGs in the human gut remains insufficiently explored. Therefore, systems level investigations are needed to determine whether antibiotic administration alters the dissemination potential of ARGs, and more importantly, which factors are associated with the altered dissemination potential of ARGs in the human gut.

In this study, we first analyzed public metagenomic data from a longitudinal study of 18 cefprozil treated and 6 control volunteers [12]. We surveyed the strain-level dynamics, shift of the resistome structure, evolution of resistance genes at the single nucleotide level, and factors associated with the variation of dissemination potential. In this first stage of *in silico* analysis we demonstrated the diversification of the resistome and *in situ* evolution of strains upon antibiotic intake. We then performed an in-depth prospective and functional study on longitudinal samples from one cefuroxime treated and one control subject using both culture-dependent and culture-independent approaches.

## Results

### Antibiotic treatment diversifies the resistome composition and increases the copy number of ARGs at the intraspecies level

We analyzed public metagenomic data from a longitudinal study of 18 cefprozil treated and 6 healthy control volunteers [12] (each subject was sampled at day 0, 7, and 90) that aimed to investigate whether the initial taxonomic composition of the gut microbiota is associated with the reshaped post-antibiotic microbiota. Using this data we investigated the dynamics and diversification of the resistome, strain-level selection, variation of the dissemination potential of antibiotic resistance, and the single-nucleotide level differentiation under antibiotic treatment.

To evaluate to what extent the composition of the resistome was altered in response to antibiotic treatment, we quantified the compositional distance between samples at different time points. An NMDS plot based on the Jaccard distance of ARGs presence/absence profile of each sample revealed that the resistome composition was more drastically altered in the antibiotic treated group than the control group during treatment (Figure 1A). The compositional distances between baseline and treatment samples within the same treated individual were significantly larger than those in the control group (0.14 *vs.* 0.06 on average, *P* value < 0.01 with Wilcoxon rank-sum test, Figure 1A). Additionally, we observed that the compositional differences between post-treatment (90 days) and the baseline samples within the same individual were also significantly larger in the treated group than in the control (0.135 *vs*. 0.075 on average, *P* value = 0.04, Wilcoxon rank-sum test, Figure 1A), revealing the persistent diversified ARG composition after antibiotic perturbation.

**Figure 1.**
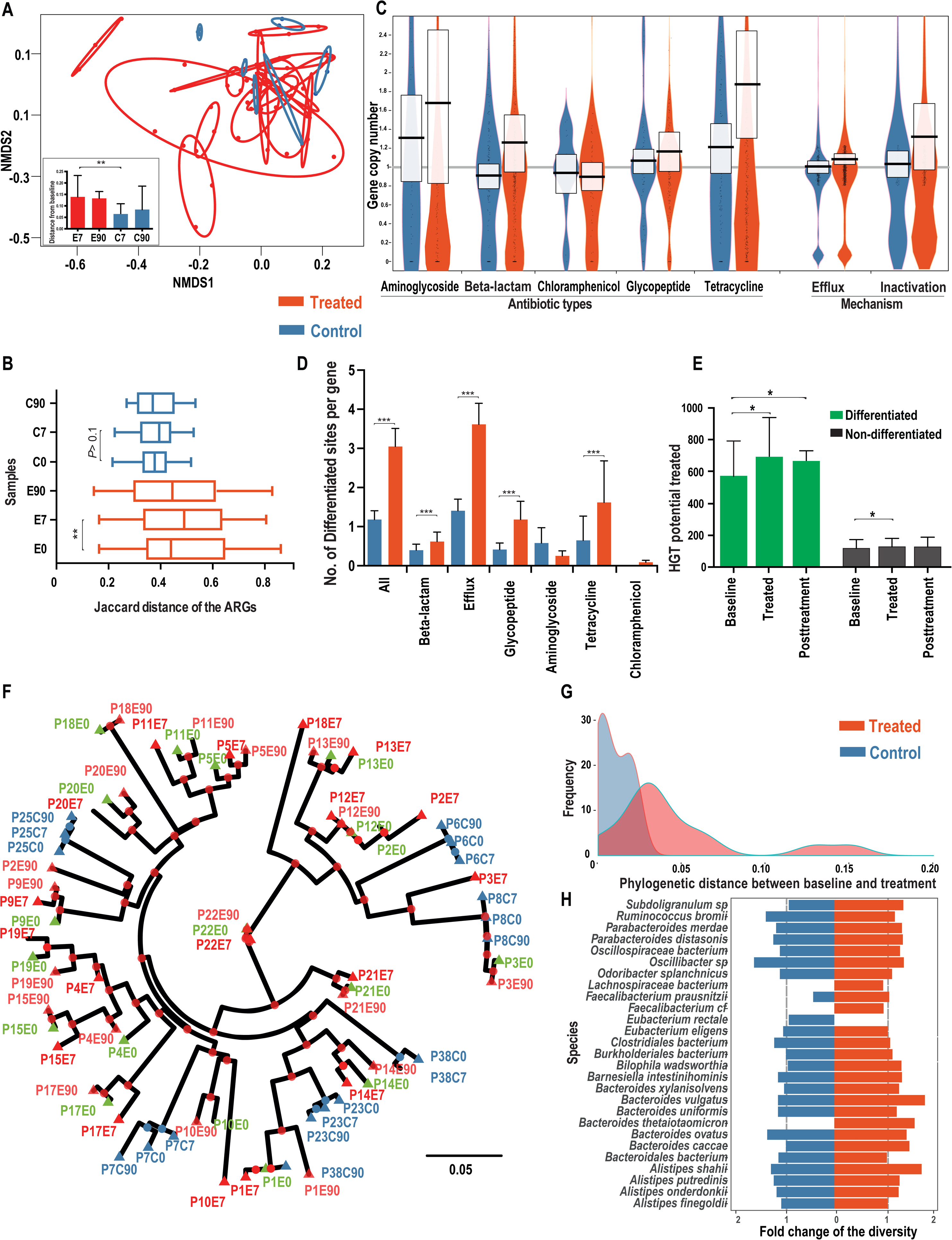
Divergence of the resistome and dominant strains in the prevalent species upon antibiotic intake. **A.** NMDS based on the Jaccard distance of antibiotic resistance gene profiles (presence/absence) (stress value < 0.05). E7, E90, C7, C90 represent the samples from Day 7 treatment, Day 90 post-treatment, Day 7 control, Day 90 control samples, respectively. The red and blue ovals surround samples from the same antibiotic treated and control subject, respectively. The radii of the ovals reflect the compositional distances between samples from the same individual. A larger radius suggests a more drastically diverged resistome profile. **B.** Distribution of the Jaccard distance of ARGs profile between individuals at the same time point. **C.** Fold-change of the ARG copy number of major categories of ARGs. The boxes describe the mean and the standard deviation. **D.** Distribution of the percentage of differentiated site in various ARG categories. **E.** Variation of the HGT potential over time for differentiated and non-differentiated ARGs, in treated subjects at baseline, treatment, and post-treatment timepoints. **F.** Phylogenomic tree based on the concatenated dominant strains sequences of 28 prevalent species. Green, red, and blue colors denote the baseline of the treated, other time points of the treated, and control samples, respectively. The samples were named using the rules of “P+individual ID+E/C (treated or control group)+time point (day 0, 7, and 90), *e.g.*, “P3E7” refers to the sample from day 7 (under antibiotic treatment) from individual 3. The green color denotes the baseline of treated subject. **G.** Density distribution of the phylogenetic distance between baseline and treated samples in Figure 1F. A larger phylogenetic distance reveals a faster evolution under antibiotic pressure. **H.** Distribution of the fold-change of the genome-wide nucleotide diversity in 28 prevalent species during the treatment. \**P* < 0.05; \*\**P* < 0.01; \*\*\**P* < 0.001

We next studied whether antibiotic treatment could select similar sets of antibiotic resistance genes and converge the resistome composition across individuals. If the gut resistome has a more common response, instead of an individualized response, to an antibiotic treatment, we would expect the resistome compositions to become more similar after the treatment across individuals. We found that the dissimilarity of the ARGs composition measured by the Jaccard distance between individuals increased significantly over time in the antibiotic treated group (baseline: 0.44 *vs*. treatment: 0.52, on average, *P* value < 0.01 with Wilcoxon rank-sum test, Figure 1B), indicating that the overall composition of the resistome diversified under antibiotic treatment across individuals. No significant increase of resistome divergence was observed in the control group (*P* value > 0.05 with Wilcoxon rank-sum test, Figure 1B). When considering the ARGs abundance rather than the presence/absence, we observed a similar pattern. The Bray-Curtis distance based on the ARG abundance profile between individuals increased significantly during treatment (0.615 *vs*. 0.685, *P* value < 0.01 with Wilcoxon rank-sum test, Figure S1). The diversified resistome suggests that antibiotic exposure drives an individualized selection of antibiotic resistance genes. Additional statistical analyses revealed that both the baseline species-level taxonomic composition and the baseline resistome composition are significantly correlated with the resistome composition after treatment (*P* values < 0.001, Mantel test with permutation = 5000).

The significantly diversified resistome composition during antibiotic treatment encouraged us to investigate whether the copy number of the ARGs (the ratio of gene depth to the relative genomic abundance of the species harboring this gene) could be altered during treatment. Incorporating the reference genomes of the 28 most prevalent species (at least 1% of relative abundance in at least half of the samples from 24 individuals, Table S1), we quantified the average copy number of each ARG. We found that genes annotated to confer resistance towards most classes of antibiotics, including aminoglycosides, beta-lactams, tetracyclines, and glycopeptides, increased the copy number significantly by 22% on average during the treatment (*P* value < 0.001, Wilcoxon rank-sum test, Figure 1C). The copy number of efflux pumps annotated as ARGs also increased significantly by 8% (*P* value < 0.001, Wilcoxon rank-sum test, Figure 1C). In contrast, ARGs in the control group had no significant increase in copy number (*P* value = 0.76, Wilcoxon rank-sum test, Figure 1C) during the same period. The significant increase in copy number of antibiotic resistance genes in response to antibiotic treatment could be explained by either the selection of strains with existing ARGs (one or multiple copies) or the horizontal transfer of ARGs.

### Antibiotic treatment drives single nucleotide level differentiation at non-synoymous sites

Next, we investigated how the genotype of ARGs varied at the single nucleotide level within each species’ population during antibiotic treatment. The analysis of single nucleotide variants (SNV) based on the shotgun metagenomic data of 72 samples from 24 individuals revealed that there is a significantly higher proportion of differentiated sites (see Methods for definition) in the treated subjects than the control group (3.0% *vs*. 1.1%, *P* value <0.001, Wilcoxon signed-rank test, Figure 1D). There is also a significantly higher proportion (31% *vs*. 13%, *P* value < 0.001 with Wilcoxon signed-rank test, Figure S2A) of genes containing differentiated sites in the treated subjects than the controls, consistent with antibiotic exposure driving compositional changes in ARG allele frequency and spectrum. By examining sub-categories of ARGs, we observed that most genes, including those annotated as efflux pumps or conferring resistance towards beta-lactams, tetracyclines, and glycopeptides, have a significantly higher proportion of differentiated sites in the treated group compared to the control (*P* value < 0.001, Wilcoxon signed-rank test, Figure 1D), indicating antibiotic-driven differentiation of the human gut resistome.

To evaluate the potential functional influence of the differentiated sites in the treated subjects, we investigated the frequency of differentiation at either non-synonymous sites (0-fold degenerate sites) or synonymous sites (4-fold degenerate sites). Compared with all the sites in the coding region, single nucleotide differentiation is drastically enriched at non-synonymous sites (91.1% *vs*. 64.9% for differentiated sites and all sites, respectively), suggesting that intraspecies level selection tended to influence the functional potential of the ARGs within the population of each bacterial species.

We further investigated whether there are some commonly differentiated sites in the ARGs among antibiotic treated individuals. We found that 11 genes habored at least one recurrent differentiated site, and these genes were observed in at least 25% of the treated individuals. These commonly differentiated genes, which belong to the MexE or RND efflux family, originate mainly from the species *Bacteroides uniformis* and *Bacteroides vulgatus*, or other *Bacteroides* species (Table S2).

### Increased HGT potential of the resistome is associated with SNV differention

We further investigated whether the dissemination potential of ARGs was altered during antibiotic treatment. The chance of HGT transfer (HGT potential) for a particular gene is associated with two factors, the intrinsic HGT tendency of the gene and its prevalence in the population. Therefore, we estimated the HGT potential for the ARG families by combining the relative abundance and the HGT rate [24] (HGT potential = HGT rate × abundance, see Methods). According to the above definition, the gene-level variation of HGT potential is noticeably correlated with the abundance variation. We found that the HGT potential increased in both differentiated and non-differentiated ARG families during antibiotic treatment (*P* value < 0.05 with Wilcoxon signed-rank test), while the HGT potential in differentiated ARGs increased more drastically and significantly than in the non-differentiated ones (19% *vs*. 5% on average, *P* value < 0.01, Wilcoxon signed-rank test, Figure 1E). This suggested that ARGs with differentiated sites could play more important roles in dissemination of antibiotic resistance upon antibiotic intake. The pattern of the differentiation-associated increase of HGT potential was not observed in the control subject at the same time period (Figure S2B).

We discovered a significantly positive correlation between community-level fold-change of the HGT potential and the average rate of differentiated sites in the ARGs (Pearson correlation coefficient 0.50, *P* value = 0.038, Figure S3B), suggesting that ARGs with a higher proportion of differentiated sites tend to increase their HGT potential more drastically. One explanation for this differentiation or SNV-associated increase of the HGT potential is that ARGs with multiple genotypes co-exist within the population for particular species before the antibiotic treatment and these genotypes confer distinct selective advantages or link with other beneficial alleles of ARGs under antibiotic pressure. Therefore, antibiotic treatment tends to select ARGs with selectively advantageous mutations, increasing their abundance and HGT potential. In constrast, non-differentiated ARGs genes harboring homogeneous alleles or multiple alleles with similar selective advantage under antibiotic pressure would randomly select the strains harboring them and maintain their allele frequency.

### Antibiotic pressure shifts the intraspecies population structure

The diversified resistome composition and single nucleotide differentiation revealed the influence of antibiotic treatment on the human gut resistome. Therefore, we further investigated how antibiotic pressure drives changes in the intraspecies-level population structure of intestinal species regarding the dominant strain genotypes. We reconstructed the phylogenomic tree for the concatenated genomes of the dominant strains from 28 prevalent species (see Methods). Our results reveal that the 28 dominant strains of the treated subjects present significantly closer phylogenetic relationships between samples from the same individual than the samples across individuals (*P* value < 0.01, permutation test, Figure 1F), indicating an individualized antibiotic selection for the dominant strains within each subject. This individual-specific strain selection during treatment was also observed for each individual species, *e.g., Ruminococcus. bromii* (Figure S4A). Furthermore, we noticed a significantly higher divergence (phylogenetic distance) of the dominant strains between the baseline and treated samples in the treated group than the control (0.052 *vs*. 0.015 on average, *P* value < 0.01, Wilcoxon signed-rank test, Figure 1F-G), suggesting that antibiotic pressure drove a significant shift of the dominant strain genotype, therefore influencing the intraspecies population structure. This is consistent with the aforementioned strong differentiation trend for the ARGs. Interestingly, when we examined the post-treatment (day 90) recovery patterns by comparing the phylogenetic distances of the dominant strains between baseline treated and post-treatment samples in the treated group, we found that the dominant strains in only 39% (7 out of 18) of the individuals had a closer phylogenetic relationship with the corresponding baseline samples than the treatment samples (Figure 1F). This suggests that the shift in intraspecies population structure upon antibiotic treatment could be long-lasting.

If antibiotic treatment tends to select dominant strains or ARGs with more similar genotypes, we should observe smaller phylogenetic distances between treatment samples across individuals, as compared with the distances between the baseline samples across individuals. The results show that there are no significantly different phylogenetic distances between the cross-individual treatment samples and cross-individual baseline samples of the dominant strains or domain ARG genotype (*P* value >0.05, Wilcoxon signed-rank test, Figure 1F and S4B), suggesting that antibiotic treatment tends to select dominant strains and ARGs with individual-specific genotypic signatures.

Furthermore, we found that the differences in genome-wide nucleotide diversity in 28 prevalent species are not significant between the treated and control groups (average fold-change: 1.12 *vs*. 1.05, *P* value = 0.73, Wilcoxon signed-rank test, Figure 1H), indicating no large-scale selective sweep or strain domination in most of the species. Interestingly, we did find in particular individuals and species (*e.g., Bacteroides uniformis* in individuals 10, 13, 17, and 19) a drastically decreased (less than half of the baseline) genome-wide nucleotide diversity (Table S3), suggesting that incomplete selective sweep occured due to antibiotic exposure. Combined with the strong divergence of the dominant strains in the treated group, our analysis indicates that cefprozil intake reshapes the intraspecies population structure by selecting individual-specific strains and ARGs.

### Cefuroxime treatment increases antibiotic resistance levels of the human gut microbiota

The analyses based on public data revealed the general tendency for differentiation and diversification of the resistome upon antibiotic treatment. Subsequently, we prospectively and functionally assessed the impact of antibiotic treatment on the resistance levels of the gut microbiota to a panel of antibiotics in one treated and control subject using both cultivation-based multiplex phenotyping and cultivation-independent functional and shotgun metagenomics (Figure 2A and Methods). We defined the level of phenotypically resistant gut microbiota as the ratio of colony counts on the antibiotic-containing plates to the antibiotic-free plates. The overall level of phenotypically resistant gut microbiota increased significantly by 140% (*P* value < 0.05, Wilcoxon rank-sum test) on the β-lactam plates during cefuroxime treatment (day 5) (Figure 2B-C, Table S4). Even though the phenotypic resistance levels to various β-lactams partially recovered after the treatment, resistance never returned to their initial levels, even 3 months after treatment. In addition, the resistance levels of non-β-lactam plates showed a moderate (49% on average) but statistically significant increase (*P* value < 0.05, Wilcoxon rank-sum test) during treatment in the cefuroxime-treated subject (Figure 2B-C), suggesting that certain bacteria, possibly harboring multiple types of antibiotic resistance genes or multiple-antibiotic resistance genes, were selected during cefuroxime treatment. In contrast, no statistically significant increase in resistance was observed in either β-lactam or non-β-lactam antibiotic plates in the control subject (Figure 2C).

**Figure 2.**
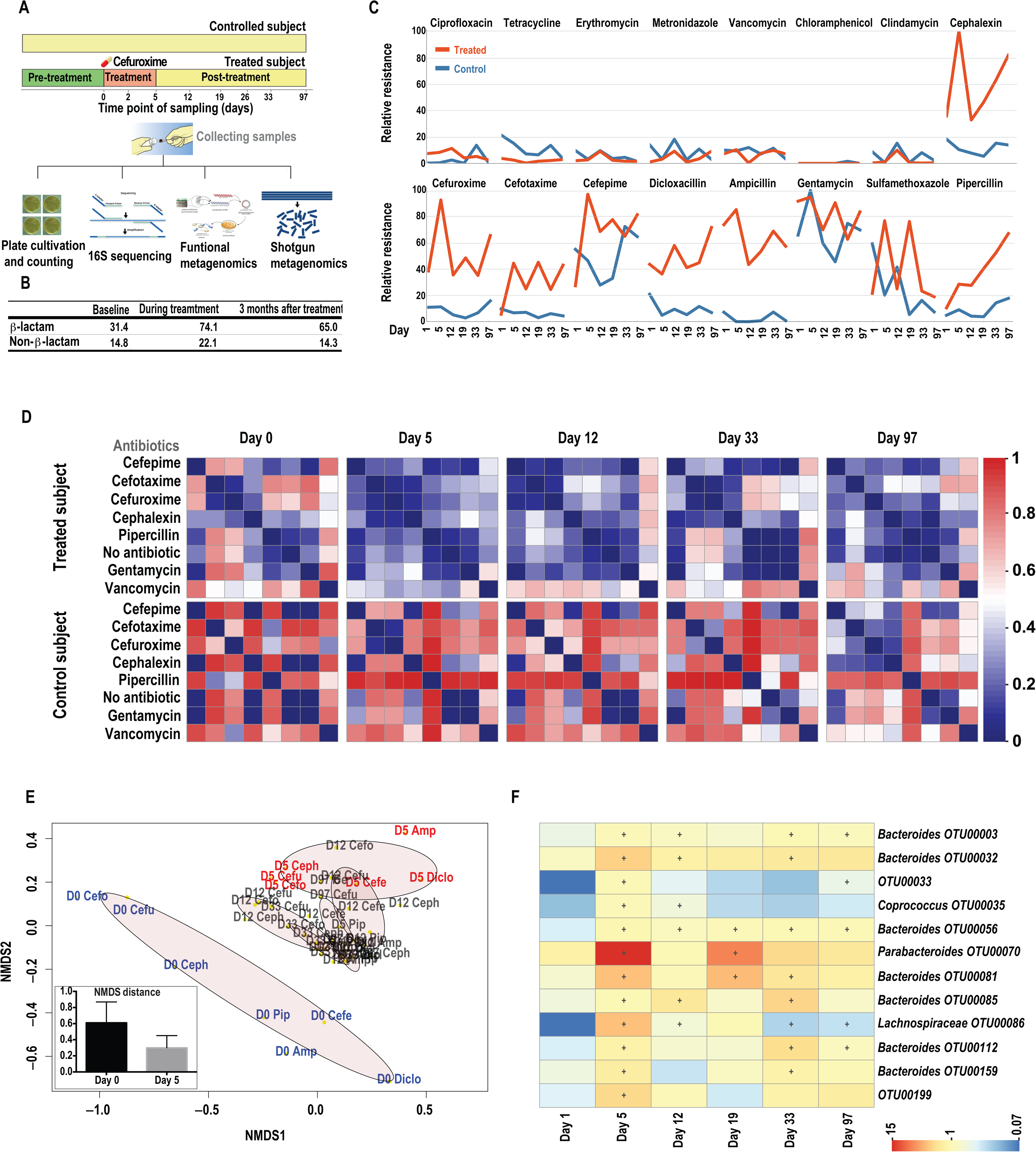
Cefuroxime treatment enhanced the resistance to multiple antibiotics and converged the cultured resistant bacterial communities. **A.** Experimental design: Two healthy adult human donors were selected for this study. One subject underwent a standard 5-day treatment with cefuroxime (500 mg, 3 times a day) while the other subject was not treated, serving as a control. Fecal samples from multiple time points were extracted for downstream experiments, including cultivation-based multiplex phenotyping, DNA sequencing of 16S rRNA, and functional and shotgun metagenomics (see Methods for more details). **B.** Summary of the average percentage resistance levels before, during, and after treatment from both β-lactam and non-β-lactam plates in the treated subject. **C.** Variation of the relative antibiotic resistance over time in the presence of different antibiotics. **D.** Heatmap of the pairwise community dissimilarity, as measured by the Morisita-Horn index, across cultivated plates at different time points in the treated and control subjects. **E.** NMDS based on Morisita-Horn dissimilarity of communities cultured with different β-lactam antibiotics in treated subjects (stress value < 0.05). The brown ovals surround the samples from sample time points. Cefo: cefotaxime, Ceph: cephalexin, Cefu: cefuroxime, Pip: pipercillin, Cefe: cefepime, Amp: ampicillin, Diclo: Dicloxacillin. **F.** Heatmap of antibiotic resistance from β-lactam plates for the species with significantly boosted resistance during treatment. + denotes the resistance is significantly higher (BH adjusted *P* value < 0.01 with permutation test) than that in the baseline sample in the treated subject.

To evaluate the variation in taxonomic composition of cultured resistant bacteria in both cefuroxime-treated and control subjects at different time points, we performed 16S rRNA sequencing of the colonies grown on different antibiotic containing plates (Table S3). We observed that the community dissimilarity (beta-diversity), measured by the Morisita-Horn index, between antibiotic plates from the treated subject decreased after the cefuroxime treatment. This converging pattern of resistant communities among different antibiotic plates was not observed in the control subject (Figure 2D and Figure S5). In addition, the pattern of converged taxonomic composition of the resistant bacteria is significantly stronger (*P* value < 0.01, Wilcoxon rank-sum test) on β-lactam plates than on the non-β-lactam plates (Figure 2D and Figure S6). An NMDS plot based on the 16S rRNA taxonomic profiles of each cultured antibiotic plate revealed that the taxonomy profiles on β-lactam plates were significantly altered (*P* value < 0.01, permutational MANOVA test) over time (Figure 2E). Such strong differentiation was not observed on the non-β-lactam plates (*P* value > 0.05, permutational MANOVA test).

By comparing the relative resistance of each species (16S based taxonomy) on β-lactam plates between baseline (day 0) and treatment (day 5) samples, we identified 12 species with significantly increased phenotypic resistance during treatment (BH adjusted *P* value < 0.05 using permutation test) from all β-lactam plates in the cefuroxime-treated subject (Figure 2F). Most of these species, mainly from the genus *Bacteroides*, sustained enhanced resistance for one to three months (Figure 2F). We further implemented the same statistical tests on non-β-lactam plates from the cefuroxime-treated subject, as well as on both β-lactam and non-β-lactam plates from the control subject. No species with significantly increased resistance were identified, suggesting that the cefuroxime treatment might select for species with higher resistance to multiple β-lactams.

### Antibiotic treatment reduces genome-wide nucleotide diversity in *Escherichia coli* and *Enterococcus faecium*

We next investigated the temporal variation of genome-wide nucleotide diversity (clonal diversity) in different species with metagenomic analysis of the cefuroxime treated and control subjects (Figure 2A and Methods). We found that *Escherichia coli* and *Enterococcus faecium* showed exceptional variation patterns of whole genome diversity compared with all other species during cefuroxime treatment (Figure 3A). We observed strong evidence for the occurrence of genome-wide selective sweeps in these two species with a sharp decrease of genome level nucleotide diversity during treatment (Figure 3A-B), indicating that particular strains with low or intermediate initial frequency were strongly selected in the face of antibiotic pressure. These strains dominated the population with clonal expansion during antibiotic treatment and reduced the heterogeneity of the population. In addition, the temporal variation patterns between the *E. coli* and *E. faecium* were quite different in terms of their post-treatment recovery mode. The genome-wide nucleotide diversity of *E. coli* recovered gradually but never returned to its initial level after the cefuroxime treatment (Figure 3A), suggesting the persistence of antibiotic selected strains of *E. coli*. This incomplete recovery of the nucleotide diversity indicates that the fitness of the strain surviving antibiotic treatment was still competitive within the population in the antibiotic-free environment. In contrast, the diversity of *E. faecium* recovered rapidly after antibiotic treatment, suggesting that the antibiotic selected strain had a fitness cost in absence of antibiotic treatment, as indicated in previous studies [25,26]. No genome-wide reduction of nucleotide diversity was observed in other species during treatment in the treated subject (Figure 3A). We observed random temporal variations of the genome-wide diversity in the control subject, which remained relatively stable over time (Figure S7), indicating that the overall strain level dynamics in the gut microbiome are more pronounced during antibiotic treatment, consistent with the observation using the public data.

**Figure 3.**
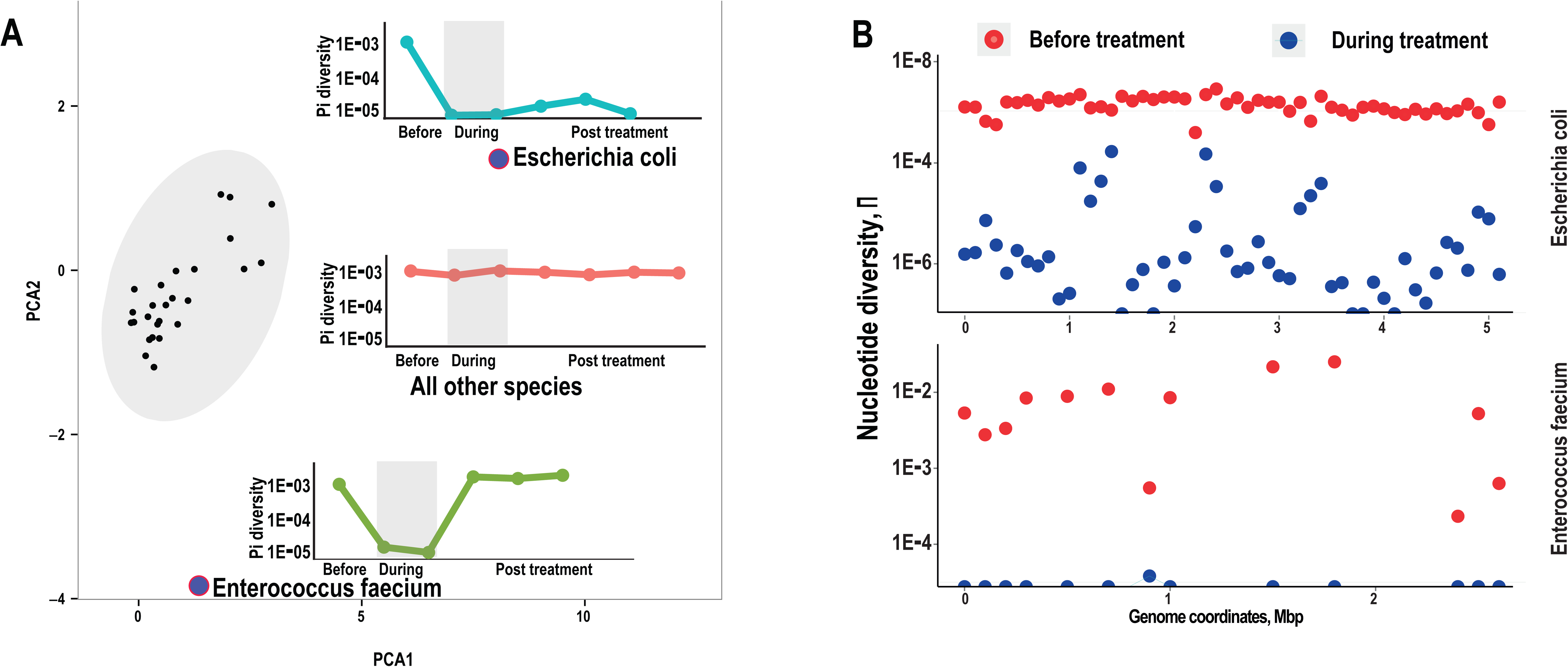
The variation of the genome-wide nucleotide diversity. **A.** PCA based on the genome-wide nucleotide diversity at different time points in the treated subject. Each dot represents one species. Two species, *Escherichia coli* and *Enterococcus faecium* show exceptional genome-wide selective sweeps. Three sub-figures describing the pattern of the temporal variation at the genome-wide diversity were shown juxtaposed with the species. **B.** The nucleotide diversity before and during treatment across the genome coordinates in *Escherichia coli* and *Enterococcus faecium* in the treated subject.

### Functionally selected ARGs experience strong selection at the single nucleotide level

We next adopted functional metagenomics to investigate the influence of the cefuroxime intake on the prevalence of different types of functionally validated ARGs over time. The results from all functional selections before treatment identified various types of ARGs, while β-lactamases increased drastically and dominated (87% of relative abundance) the recovered genes (Figure 4A-B) during the cefuroxime treatment. The β-lactamases decreased post-treatment but remained prevalent (37%) in the functional selections (Figure 4A). When quantifying the ARGs abundance from β-lactam and non-β-lactam plates separately, we noticed a very similar dominance of β-lactamases on the β-lactam plates during the treatment (Figure S8A). Additionally, we detected an increase of efflux genes on the non-β-lactam plates during the cefuroxime treatment period (Figure S8B), suggesting co-selection of other resistance genes. This consistent with our results from the cultivation plates, where resistance levels were also enhanced on some non-β-lactam antibiotic plates. To validate the increased abundance of β-lactamases in the gut microbiota of the treated subject, we mapped the shotgun metagenomic data to the functionally validated β-lactamases. The results showed that β-lactamases increased drastically during treatment in the cefuroxime-treated subject while there was no such trend in the control subject (Figure S9).

**Figure 4.**
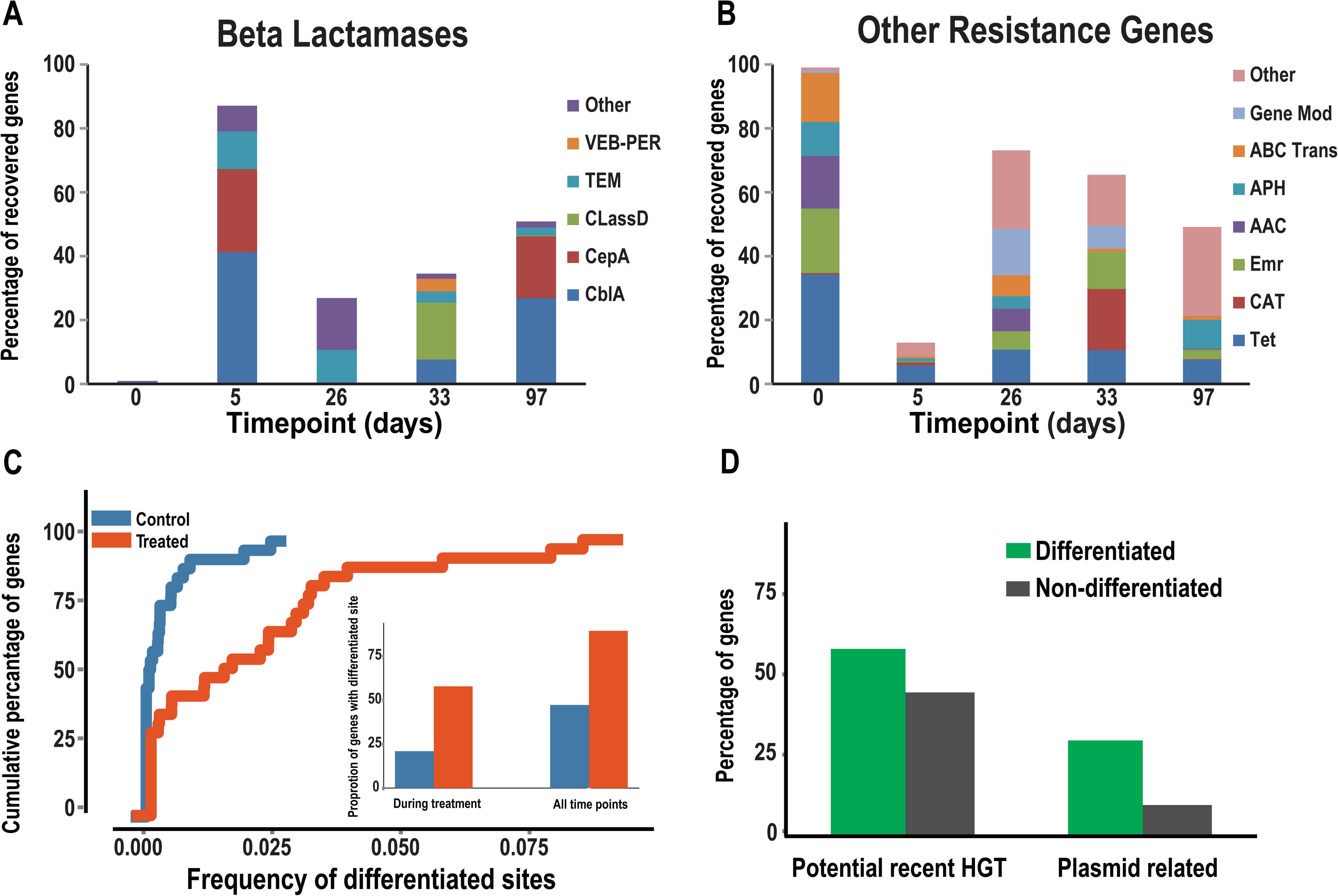
Variation of the gene abundance, differentiated sites, and HGT trend in the functional metagenomic ARGs. **A.** Temporal variation of the gene abundance for functionally selected β-lactamases. **B.** Temporal variation of the gene abundance for functionally selected non-β-lactamase antibiotic resistance genes. **C.** Cumulative percentage of functional ARGs with different proportions of differentiated sites. The proportions of genes with differentiated sites before and during the treatment in both subjects are shown in the subfigure. **D.** Proportion of genes with recent HGT signatures and genes with plasmid mediated HGT in both differentiated and non-differentiated ARGs.

Since the genome-wide selective sweep in *E. coli* and *E. faecium* revealed a strong selective force imposed by antibiotic treatment at the intraspecies level, we subsequently investigated further how the functional ARGs diversified within the population of each species during the cefuroxime treatment. The analysis of SNV, combining functional and shotgun metagenomics data, revealed that the 114 functionally selected ARGs contain a significantly higher proportion of differentiated sites in the cefuroxime-treated subject than in the control (0.037 *vs*. 0.004, *P* value < 0.01 with Wilcoxon signed-rank test) (Figure 4C). At the same time, the proportion of functional ARGs with SNV signals of selection (genes with at least one differentiated site) is significantly higher in the treated subject than in the control (56% *vs*. 22%, *P* value < 0.01 with Fisher’s exact test) during or after treatment (Figure 4C), consistent with our findings based on the computationally annotated metagenomics datasets (Figure 1D). Further analysis revealed that the nucleotide diversity (π) of ARGs decreased significantly (*P* value < 0.01 with Wilcoxon signed-rank test) during treatment in the cefuroxime-treated subject (Figure S10), consistent with the strong diversification of the allele frequency at the single nuecleotide level.

### Differentiated functional ARGs tend to be recently horizontally transferred

To evaluate the taxonomic origin and potential of recent HGT in functionally selected ARGs, we identified the last common ancestor (LCA) of highly homologous hits (identity >95% at amino acid level) of the functional ARGs against the NR database using blast (see Methods). We explicitly defined recent HGT-related ARGs by considering the confidence level of LCA and the taxonomic information from NR hits (see Methods). This analysis indicates that there is a higher proportion of differentiated functionally selected ARGs involved in recent HGT events than in the non-differentiated ones (57% *vs*. 42%) (Figure 4D). To validate this trend of recent HGT in differentiated ARGs, we incorporated the public data described above [12] and observed a very similar pattern— the differentiated ARGs tend to be more recently transferred than the non-differentiated ones (Figure S11), suggesting a stronger HGT trend in the differentiated ARGs.

The higher rate of recent HGTs in differentiated ARGs motivated us to investigate further whether these ARGs were associated with plasmid-mediated bacterial conjugation. We searched for all highly homologous hits (> 95% identity) in the NCBI plasmid RefSeq database using the ARGs as queries. We found that about 33% of differentiated and 11% of non-differentiated ARGs have highly similar (> 95% identity at amino acid level) plasmid hits (Figure 4D), supporting the role of plasmids in harboring and circulating these ARGs. Since only plasmid hits with high identity (> 95%) were considered, we can be quite confident that a substantial proportion (33%) of differentiated ARGs are actively transferred among species via plasmids in the recent evolutionary history. In our prospective study, we also observed that the HGT potential in differentiated ARGs increased more drastically than in the non-differentiated ones (304% *vs*. 142%, *P* value < 0.01 with Wilcoxon signed-rank test, Figure S12) as we proposed above using the public dataset.

## Discussion

In this study, we observed antibiotic-induced strain-level dynamics, resistome diversification, and increased resistance dissemination potential within the human gut. The integrative approach deployed in this study enabled the elucidation of complex dynamics of the resistome and gut microbial strains during antibiotic treatment. We used computational analyses of existing metagenomic datasets to find evidence for antibiotic induced diversification of the resistome, as well as substantial treatment induced differentiation of antibiotic resistance genes. We further elucidated widespread strain level dynamics exacerbated by antibiotic treatment. Combining these results, we showed that the increased dissemination potential of antibiotic resistance gene is strongly associated with the frequency of the differentiated sites during antibiotic treatment. We then performed a functional survey in a prospective intervention study, enabling detailed culture-based and culture-independent characterization of the human gut resistome and resistance phenotypes. The consistent findings from two experimental designs reflect the generalization of our conclusions.

Previous studies have provided evidence that certain factors could influence the HGT potential in the human intestine. For instance, it has been shown that human intestinal epithelial cells produce proteinaceous compounds that modulate antibiotic resistance transfer via plasmid conjugation in *E. coli* [27]. Another study using a mouse colitis model demonstrated that pathogen–driven inflammation of the gut promoted conjugative gene transfer between *Enterobacteriaceae* species due to the transient bloom of the pathogenic *Enterobacteriaceae* [28]. Our results revealed a similar but more comprehensive scenario about the increased HGT potential under antibiotic treatment, supporting the hypothesis that antibiotic pressure could drive the dissemination of the resistome [16]. According to the definition of HGT potential, the overall increase of ARGs abundance could be the major reason for the overall increased HGT potential, although the increased ARG abundance may neither guarantee an increased HGT potential (see Methods), nor is it proportionate to overall increased abundance [24]. More importantly, we observed that the increased HGT potential of ARGs was compounded by antibiotic selection of these genes at the single nucleotide level, highlighting the association between evolutionary plasticity and the horizontal transfer of ARGs [29]. The single nucleotide level differentiation could be explained by the existence of a huge multiplicity of bacterial clones within single species in the gut before the antibiotic perturbation. Antibiotic treatment is the driving force selecting the alleles conferring higher survival advantage, leading to genotype differention and abundance increase.

Due to the fact that the disturbed microbiota could lead to adverse health outcomes [30], secondary infections [31], or increased risk of colorectal cancer [32], personalized medicine for a bacterial infection should in the future incorporate information of the patient’s gut microbiota. The knowledge of the personalized gut microbiota sets the basis for predicting the stability or dynamics at the whole community level or individual bacterium upon perturbation [33]. Although we cannot predict the clinical impact or the gut microbe dynamics due to insufficient sample size and lack of clinical records in our study, our data suggests that antibiotic therapy leads to personalized resistome diversification and individual-specific, strain-level selection in the gut microbiota. The selective sweep observed in *E. coli* and *E. faecium* highlights the influence of antibiotic treatment on the intraspecies level dynamics. In line with this, future antibiotic treatment should be more personalized regarding the dosage, duration, and combination of drugs based on the unique strains and resistome composition in each patient to minimize the unintended disruption of the gut microbiota. Unfortunately, how the initial gut microbiota composition relates to antibiotic treatment efficacy, side effects, long-term susceptibility to different pathogens, or diseases is poorly explored thus far. Therefore, more studies with longitudinal sampling and sequencing of the gut microbiota, evaluation of antibiotic efficacy, and survilliance of susceptibility for infections or other diseases are needed. Such data could uncover the microbiota-dependent antibiotic efficacy and side effects, the interaction networks between antibiotic and gut microbes, and long-term microbial dynamics, paving the way for future microbiome-based diagnosis and treatment.

Although our strain-level analyses offer novel insights into the dynamics of the resistome compostion, copy number variation, and single nucleotide level differentiation of ARGs upon antibiotic treatment, the scarcity of reference strain genomes for many species and the incompeleteness of the ARG database as well as the non-robust annotation methods could potentially bias these quantifications to certain extent and limit the generalization of our conclusions. Another caveat is that although metadata, including gender, age, weight, etc., for each individual were provided in the original study of the public data, the sample size was not sufficient for further in-depth analyses regarding these potentially confounding factors. Future studies with more individuals, increased sampling density, and the development of more comprehensive ARGs databases and accurate annotation methodologies would be of great value.

## Materials and methods

### Retrieval of public data

A total of 72 samples from 18 cefprozil treated subjects and 6 control subjects used in the study of Raymond et al. [34] were downloaded from NCBI SRA PRJEB8094. Each subject provided three longitudinal samples—baseline (day 0), treatment (day 7), and post treatment (day 90).

### Experimental Design

To functionally validate the findings of our computational analysis based on public metagenomic datasets, two healthy adult human female subjects (age 25-29, diet not controlled) who had not taken any antibiotics for at least one year were selected for this study. One subject underwent a standard 5-day treatment with cefuroxime (500 mg, 3 times a day) while the other subject had no treatment, serving as a control. Eight fecal samples were collected longitudinally over a period of three months corresponding to pre-treatment (Day 0), two time points during treatment (Day 2 and 5), one week post-treatment (Day 12), two weeks post-treatment (Day 19), three weeks post-treatment (Day 26), one month post-treatment (Day 33), and three months post-treatment (Day 97). All participants consented to these experiments and sample collections and downstream experiments and data processing followed ethical guidelines (Hvidovre Hospital) throughout the study. Samples were transported to an anaerobic chamber within an hour of collection. Five grams were separated out for culturing (only on Day 0, Day 5, Day 12, Day 19, Day 33, and Day 97) and 2.5 grams were used for DNA extractions. The remaining stool samples were stored at −80°C.

### Cultivation, DNA extraction, and 16S gene sequencing

The stool samples were cultivated with or without the presence of 16 antibiotics, followed by DNA extraction and sequencing of 16S rRNA as described previously [35]. Briefly, five grams of fecal sample was resuspended in 50 mL of prereduced (resazurin, 0.1 mg/mL) 1× PBS and 10-fold serial dilutions were plated on Gifu Anaerobic Media (GAM) agar with or without antibiotics in duplicate. The plates were incubated anaerobically at 37°C for 5 days. Bacterial colonies were then manually scraped off the surface of the agar and collected in 10 mL tubes. DNA was extracted from the collected samples using the MoBio UltraClean Microbial DNA Isolation kit.

### Identification of the species with enhanced resistance from cultured plates

The relative resistance level for each OTU is defined as the ratio of the relative abundance of the OTU from each antibiotic plate to the relative abundance of the same OTU in the control plate at each time point. To test whether certain OTUs had overall enhanced resistance over time, the resistance levels (*e.g.*, all β-lactam plates) for each OTU were compared using a Wilcoxon signed-rank test [37] between the test time points (during or post treatment) and the baseline.

### DNA extraction, library construction, and sequencing for culture-independent methods

The fecal samples for culture-independent methodologies were extracted using 2.5 grams of sample with the MoBio PowerMax Mega Soil DNA Isolation kit following the standard protocol. DNA from the treated subject’s samples from Day 0, Day 5, Day 19, Day 26, and Day 33 was used to construct functional metagenomic libraries as modified from Sommer et. al. [19].

All DNA fragments from 1056 functional clones were sequenced using Sanger sequencing, imported into CLC Main Workbench where the cloning vector sequence was removed and reads of poor quality were discarded. Assembly of reads from all samples was attempted simultaneously in order to simplify the final output and identify sequences sharing the same sequence. All assembled contigs were hand checked for errors and correct alignment. A total of 197 contigs consisting of 2 or more sequences and 104 unassembled single sequences were retrieved. Open reading frames (ORFs) were annotated using ORFinder (http://www.ncbi.nlm.nih.gov/gorf/gorf.html). ORFs were annotated by comparing the protein (pFAM database, blast×, cdd database) or nucleotide (blastn, CARD database, ARDB database) sequences to known sequences in several databases with 80% identity and coverage. Sequences reads were removed from the annotation list if no match to known or suspected antibiotic resistance genes could be found. Forward and reverse reads from the same sequence that overlapped along the same resistance gene(s) were merged into a single annotation.

### Deduction of HGTs and calculation of HGT rate

The protein sequences of the functional ARGs were mapped to the NCBI NR database using blastp (−e 1E–5) first. The latest common ancestor (LCA) at species level based on highly homologous (identity > 95%) hits for each ARG was deduced using MEGAN [38]. By definition, 100% confidence of the LCA at species level reflects an explicit origin of species and 0% confidence reveals a, theoretically, infinite gene flow among species. An ARG was deduced as recent HGT related when all following criteria were satisfied, 1) more than 2 highly homologous hits (identity > 95%); 2) confidence of LCA at species level less than 50%; and 3) the highly homologous hits were observed in at least two species, excluding the hits with incomplete taxonomic information at species level. To estimate the plasmid-mediated recent HGTs, we mapped all the ARGs against the NCBI plasmid RefSeq database [39] and only hits with > 95% identity remained for further analysis.

A total of 154,805 gene families were retrieved from the HGTree database [40]. Protein sequences of the functional ARGs were mapped to HGTree families with blastp e-value 1E–5. The HGTree family with the highest number of hits, which satisfied the blast evalue < 1E–5 and coverage >50% in the short sequence of query and subject, was defined as the gene family of that ARG. Phylogenetic reconciliation analysis was carried out using RANGER-DTL [41] based on the species tree and gene tree deduced by the HGTtree. Only the interspecies level HGTs remained for downstream analysis. The number of HGTs was divided by the total length of the phylogenetic tree in this family to deduce the family-level HGT rate.

### Shotgun metagenome library construction and sequencing

Culture-independent fecal extracts from treated subject samples Day 0, Day 2, Day 5, Day 12, Day 19, Day 26, Day 33, and Day 97 and control subject samples Day 0, Day 2, Day 19, and Day 97 were used to build shotgun metagenomic libraries using the Nextera ×T kit with the standard protocol. The HiSeq 1500 was used for 100 bp PE sequencing in the CGS of The University of Hong Kong and the average throughput for each sample was 10.5 Gbp. The raw sequences can be found in BGID (CRA000815).

### Quality control for the raw sequences of shotgun metagenomic data

To retrieve the high quality reads for downstream analyses, we used a series of quality control steps to remove the low quality reads/bases as described previously [42]. In the first step, all the Illumina primer/adaptor/linker sequences were removed from each read. Subsequently, we mapped all the reads to the human genome with BWA version 0.7.4-r385 [43], and reads with > 95% identity and 90% coverage to the human genome were removed as human DNA contamination. We further filtered the low quality regions, reads, and PCR duplicates using a previously described procedure [44].

### Reference mapping, gene copy number calculation, variant calling, dominant strain identification, and annotation of antibiotic resistance genes

The Metagenomic Intra-Species Diversity Analysis System (MIDAS) [45] was adopted to calculate the gene copy number and call single nucleotide variants within each species using the default setting. To filter out low quality SNVs, the read error was controlled by a base quality score 30 and mapping quality were controlled by MAPQ 20 from MIDAS. The relative abundances of genes in each sample were further estimated using a RPKM measurement (number of reads per kbp length of gene per million mappable reads) using in-house scripts. The 28 prevalent species in the public shotgun metagenomic data were defined as the species with at least 1% of relative abundance in at least half of the samples from 24 individuals (Table S1). To annotate the antibiotic resistance genes, a hidden Markov Model based profile searching was carried out using Resfams [46] with default parameters on the pangenome genes from MIDAS. To identify the genotypes of the dominant strain of each prevalent species at either genome level or gene level, we used the script of “call_consensus.py” from MIDAS package. The phylogenetic tree was reconstructed using FastTree [47] with maximum likelihood method with default parameters.

### Definition of HGT potential

For a single gene *i*, the HGT potential (*HGTP*_*i*_) is defined as,

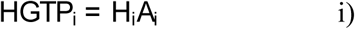

where *H*_*i*_ is the aforementioned HGT rate in this gene family *i* and *A*_*i*_ is the relative abundance of this gene in the sample.

For a set of *n* genes, the overall HGT potential is defined as,

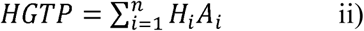

According to formula i), the increase of the HGTP for a gene is proportional to the increase of the abundance. However, for a set of genes, the overall increased abundance may not necessarily lead to an increase of overall HGTP according to formula ii). For example, a set of genes A, B, and C have HGT rates 0.5, 1, and 2, respectively. The initial and final relative abundance for these three genes are (0.1, 0.3), (0.2, 0.3), (0.3, 0.1), respectively. The overall fold-change of the abundance and HGTP is 16.7% and −23.5%, respectively. This example illustrates that the variation of the HGTP for a gene set is not only correlated to the overall abundance variation (an increased overall abundance may lead to a decreased overall HGT potential), but also the dependency between the abundance change and the HGT rate of each gene.

### Detecting differentiated sites

To identify the potential adaptively evolved variant sites in the genes, we calculated the difference of the allele frequency spectrum, which was extracted from the variants calling using MIDAS mentioned above, using Fisher’s exact test between treatment and baseline samples at each single nucleotide variant site. For example, the A:T ratio for a particular variant site in one ARG is 1:9 initially but was altered to 7:3 during the treatment, indicating potentially adaptive evolution where the allele frequency distribution has been differentiated. To correct for the influence of sequence depth on the statistical power, the maximum depth for each site was down-sampled a maximum of 10 before Fisher’s exact test was carried out. The raw *P* values were adjusted to false discovery rate (FDR) using Benjamin’s method [48].

### Statistical Analysis

The enhanced overall resistance level in the culture-dependent plates and the species with enriched resistance across multiple plates were analyzed using the Wilcoxon signed-rank test [37]. Detection of differentiated sites according to the allele frequency spectrum and the higher proportion of differentiated sites in the ARGs in the treated subjects were carried out based on Fisher’s exact test and Wilcoxon signed-rank test, respectively. The difference between resistant bacterial communities when comparing baseline and treatment plates was carried out using the permutational MANOVA test using VEGAN [49]. The NMDS analysis and the stress value calculation were performed using VEGAN. The statistical differences between groups regarding compositional distances (Bray–Curtis dissimilarity) of gut microbiota or resistome, the Jaccard distance of resistome, the copy number of ARG families, HGT potential, phylogenetic distances, genome-wide nucleotide diversity, or nucleotide diversity of ARGs, were tested using Wilcoxon signed-rank test. The statistical correlation of the taxonomic composition and resistome profile was performed by Mentel test.

## Supporting information

Information of 28 prevalent species used in the strain-level analysis.

Information of the genes with common differentiated sites across treated individuals in 28 species

The fold-change (<0.5) of genome-wide diversity during treatment for particular species.

Average colonies grown on different cultivated antibiotic plates from different time points.

The average Bray-Curtis distance based on the ARGs abundance profile between individuals.

Variation of the HGT potential over time for two types of ARGs in the control subjects.

Correlation between the variation of HGT potentials and HGT or SNP rate.

The phylogenomic tree of A. dominant strain in species Ruminococcus bromii and B. all dominant genotypes of antibiotic resistance genes.

The average Morisita-Horn dissimilarity of the resistant and controlled communities cultured.

The pairwise Morisita-Horn dissimilarity (beta-diversity) between cultured communities with antibiotics from all time points.

Variation of the genome-wide diversity over time in the control subject.

Abundance variation for various types of antibiotic resistant genes in A. beta-lactam plates and B. non-beta-lactam plates.

Abundance variation for beta-lactamases.

Temporal variation of the nucleotide diversity in the predicted ARGs in both treated and control subject.

The percentage of genes with recent HGT signatures based on public database.

Variation of the relative abundance and HGT potential of the functionally selected ARGs.

## Authors’ contributions

MOAS, EAR, and GP designed this study. EAR performed the experiments. JL, EAR, ME, and EVDH analyzed the data. JL and EAR drafted the manuscript. All authors commented on and revised the manuscript.

## Acknowledgements

This project was supported by the Lundbeck Foundatation and EU FP7-Health Program Evotar (282004). The study was approved (REG-026-2014) by the regional ethics committee and Danish national medicine agency. JL and GP would like to thank the Centre for Genomic Sciences (CGS) of The University of Hong Kong (HKU) for their support. GP would like to thank Deutsche Forschungsgemeinschaft (DFG) CRC/Transregio 124 ‘Pathogenic fungi and their human host: Networks of interaction’, subproject B5. We would especially like to thank Dr Agnes Chan (CGS) for fruitful discussions. We would like to thank Dr. Wendy Kwok for her language editing.

We thank David Westergaard for his involvement in the initial stage of the project providing the ARG annotation pipeline of the shotgun metagenomics analysis. The raw sequences were deposited in BGID (CRA000815).

## Supplementary material

**Table S1**. Information of 28 prevalent species used in the strain-level analysis

**Table S2**. Information of the genes with common differentiated sites across treated individuals (frequency of the recurrent differentiated site >25%) in 28 prevalent species.

**Table S3**. The fold-change (<0.5) of genome-wide diversity during treatment for particular species.

**Table S4**. Average colonies grown on different cultivated antibiotic plates from different time points.

**Figure S1**. The average Bray-Curtis distance based on the ARGs abundance profile between individuals.

E0, E7 and E90 refer to day 0, 7 and 90 in the treated group, respectively. C0, C7 and C90 refer to day 0, 7 and 90 in the control group, respectively.

**Figure S2**. Variation of the HGT potential over time for two types of ARGs in the control subjects.

**Figure S3**. Correlation between the variation of HGT potentials and HGT or SNP rate. **A.** Pearson correlation (correlation coefficient 0.35, P value<0.001) between the gene-level change of HGT potential and the gene HGT rate. **B.** Pearson correlation (correlation coefficient 0.50 with P value=0.038) between the gene-level change of HGT potential and the gene SNP rate (proportions of differentiated sites). The bar height is the mean value and the error bar reflect the standard deviation. The ribbons reflect the standard error of the regression. **C.** The variation of the HGT potential in 28 prevalent species across all 18 treated subjects.

**Figure S4**. The phylogenomic tree of **A.** dominant strain in species *Ruminococcus bromii* and **B.** all dominant genotypes of antibiotic resistance genes.

**Figure S5**. The average Morisita-Horn dissimilarity of the resistant and controlled communities cultured.

**Figure S6**. The pairwise Morisita-Horn dissimilarity (beta-diversity) between cultured communities with antibiotics from all time points.

Green squares surround all pairwise community dissimilarity between all plates at each time point. Red squares surround all pairwise community dissimilarity between beta-lactam plates at each time point. Cefo: cefotaxime, Ceph: cephalexin, Cefu: cefuroxime, Pip: pipercillin, Cefe: cefepime, Amp: ampicillin, Diclo: Dicloxacillin, Cipro: Ciprofloxacin, Sulfa: Sulfamethoxazole, Tet: Tetracycline, Clinda: Clindamycin, Ery: Erythromycin, Metro: Metronidazole, Vanc: Vancomycin

**Figure S7**. Variation of the genome-wide diversity over time in the control subject.

**Figure S8**. Abundance variation for various types of antibiotic resistant genes in **A.** beta-lactam plates and **B.** non-beta-lactam plates.

**Figure S9**. Abundance variation for β-lactamases.

**Figure S10.** Temporal variation of the nucleotide diversity in the predicted ARGs in both treated and control subject.

**Figure S11.** The percentage of genes with recent HGT signatures based on public database.

**Figure S12**. Variation of the relative abundance and HGT potential of the functionally selected ARGs. Left panel: relative abundance of the functional ARGs over time in both subjects. Right panel: variation of the HGT potential over time for two types of ARGs in both treated and control subjects.

