## Supplementary material for "Antibiotic Treatment Drives the Diversification of the Human Gut Resistome": Information of 28 prevalent species used in the strain-level analysis.

| **ID of species group** | **Total pangenome genes** | **# of ARGs** |
| --- | --- | --- |
| *Alistipes finegoldii* | 4,572 | 6 |
| *Alistipes onderdonkii* | 4,393 | 9 |
| *Alistipes putredinis* | 2,372 | 3 |
| *Alistipes shahii* | 3,299 | 5 |
| *Bacteroidales bacterium 58650* | 2,848 | 5 |
| *Bacteroides caccae* | 5,492 | 9 |
| *Bacteroides ovatus* | 26,316 | 42 |
| *Bacteroides rodentium* | 5,135 | 9 |
| *Bacteroides thetaiotaomicron* | 12,013 | 26 |
| *Bacteroides uniformis* | 12,228 | 28 |
| *Bacteroides vulgatus* | 22,080 | 37 |
| *Bacteroides xylanisolvens* | 16,154 | 32 |
| *Barnesiella intestinihominis* | 3,030 | 7 |
| *Bilophila wadsworthia* | 9,156 | 23 |
| *Blautia wexlerae* | 8,586 | 18 |
| *Burkholderiales bacterium* | 3,287 | 8 |
| *Clostridiales bacterium* | 13,088 | 30 |
| *Eubacterium eligens* | 2,665 | 10 |
| *Eubacterium rectale* | 7,295 | 21 |
| *Faecalibacterium cf 62236* | 2,825 | 7 |
| *Faecalibacterium prausnitzii* | 5,170 | 14 |
| *Oscillibacter sp 60799* | 2,830 | 6 |
| *Oscillospiraceae bacterium* | 5,452 | 14 |
| *Parabacteroides distasonis* | 18,432 | 40 |
| *Parabacteroides merdae* | 6,987 | 11 |
| *Ruminococcus bromii* | 2,204 | 7 |
| *Subdoligranulum sp 62068* | 4,483 | 16 |
