## Supplementary material for "Antibiotic Treatment Drives the Diversification of the Human Gut Resistome": Information of the genes with common differentiated sites across treated individuals in 28 species

| **Gene ID** | **Rasfam ID** | **Gene symbol** | **Species** | **No. of recurrent differentiated sites** |
| --- | --- | --- | --- | --- |
| 1499682.3.peg.1898 | RF0098 | MexE | *Alistipes onderdonkii* | 5 |
| 411901.7.peg.2138 | RF0098 | MexE | *Bacteroides caccae* | 13 |
| 469590.5.peg.5635 | RF0098 | MexE | *Bacteroides ovatus* | 44 |
| 411479.10.peg.1562 | RF0098 | MexE | *Bacteroides uniformis* | 2 |
| 411479.10.peg.2111 | RF0098 | MexE | *Bacteroides uniformis* | 1 |
| 997889.3.peg.2198 | RF0099 | MexH | *Bacteroides uniformis* | 5 |
| 411479.10.peg.1561 | RF0115 | RND efflux | *Bacteroides uniformis* | 8 |
| 1339352.3.peg.2493 | RF0007 | ABC efflux | *Bacteroides vulgatus* | 2 |
| 469593.3.peg.2930 | RF0115 | RND efflux | *Bacteroides vulgatus* | 1 |
| 1339350.3.peg.2181 | RF0115 | RND efflux | *Bacteroides vulgatus* | 1 |
| 742738.3.peg.4077 | RF0007 | ABC efflux | *Clostridiales bacterium* | 1 |
