## Supplementary material for "Antibiotic Treatment Drives the Diversification of the Human Gut Resistome": The fold-change (<0.5) of genome-wide diversity during treatment for particular species.

| **Individual ID** | **Fold-change of genome-wide diversity during treatment** | **Treated (T) or Control (C)** | **Species** |
| --- | --- | --- | --- |
| P18 | 0.34 | T | *Alistipes onderdonkii* |
| P10 | 0.35 | T | *Bacteroidales bacterium* |
| P10 | 0.3 | T | *Bacteroides caccae* |
| P19 | 0.33 | T | *Bacteroides uniformis* |
| P10 | 0.42 | T | *Bacteroides uniformis* |
| P17 | 0.41 | T | *Bacteroides uniformis* |
| P13 | 0.43 | T | *Bacteroides uniformis* |
| P17 | 0.4 | T | *Bacteroides vulgatus* |
| P10 | 0.26 | T | *Bacteroides vulgatus* |
| P10 | 0.47 | T | *Bacteroides xylanisolvens* |
| P15 | 0.49 | T | *Bacteroides xylanisolvens* |
| P22 | 0.38 | T | *Burkholderiales bacterium* |
| P10 | 0.25 | T | *Faecalibacterium prausnitzii* |
| P38 | 0.45 | C | *Faecalibacterium prausnitzii* |
| P9 | 0.48 | T | *Oscillospiraceae bacterium* |
| P10 | 0.49 | T | *Parabacteroides merdae* |
