## Supplementary material for "Antibiotic Treatment Drives the Diversification of the Human Gut Resistome": Average colonies grown on different cultivated antibiotic plates from different time points.

| **Control Subject** | **D0** | **D5** | **D12** | **D19** | **D33** | **D97** |
| --- | --- | --- | --- | --- | --- | --- |
| Ciprofloxacin (16 μg/mL) | 2.15E+07 | 4.35E+07 | 1.46E+08 | 2.10E+07 | 8.65E+08 | 1.25E+07 |
| Tetracycline (32 μg/mL) | 2.28E+09 | 1.50E+09 | 4.10E+08 | 1.16E+09 | 8.60E+08 | 1.34E+08 |
| Erythromycin (32 μg/mL) | 1.07E+09 | 3.49E+08 | 5.70E+08 | 7.20E+08 | 3.10E+08 | 7.35E+07 |
| Metronidazole (16 μg/mL) | 1.50E+09 | 3.07E+08 | 1.04E+09 | 4.95E+08 | 6.85E+08 | 1.19E+08 |
| Vancomycin (32 μg/mL) | 1.08E+09 | 9.25E+08 | 6.85E+08 | 1.27E+09 | 7.25E+08 | 1.31E+08 |
| Chloramphenicol (32 μg/mL) | 4.10E+04 | 1.76E+05 | 1.20E+04 | 1.70E+05 | 1.00E+08 | 6.10E+04 |
| Clindamycin (16 μg/mL) | 9.35E+08 | 6.20E+07 | 8.70E+08 | 9.70E+07 | 5.10E+08 | 9.00E+07 |
| Cephalexin (32 μg/mL) | 1.95E+09 | 1.03E+09 | 4.45E+08 | 9.65E+08 | 9.65E+08 | 5.50E+08 |
| Cefuroxime (32 μg/mL) | 1.18E+09 | 1.12E+09 | 3.05E+08 | 5.40E+08 | 4.35E+08 | 6.30E+08 |
| Cefotaxime (16 μg/mL) | 1.05E+09 | 6.75E+08 | 4.15E+08 | 5.70E+08 | 3.95E+08 | 1.81E+08 |
| Cefepime (16 μg/mL) | 5.85E+09 | 4.55E+09 | 1.59E+09 | 5.90E+09 | 4.55E+09 | 2.55E+09 |
| Dicloxacillin (32 μg/mL) | 2.23E+09 | 4.95E+08 | 5.50E+08 | 9.80E+08 | 7.40E+08 | 2.80E+08 |
| Ampicillin (32 μg/mL) | 7.85E+08 | 2.31E+07 | 1.10E+07 | 1.60E+08 | 4.70E+08 | 4.00E+07 |
| Gentamycin (32 μg/mL) | 7.00E+09 | 9.75E+09 | 3.40E+09 | 8.15E+09 | 4.70E+09 | 2.75E+09 |
| Sulfamethoxazole (64 μg/mL) | 6.30E+09 | 1.99E+09 | 2.37E+09 | 1.00E+09 | 1.03E+09 | 3.00E+08 |
| Pipercillin (32 μg/mL) | 5.40E+08 | 9.05E+08 | 2.50E+08 | 7.05E+08 | 9.10E+08 | 7.05E+08 |
| No Antibiotic | 1.07E+10 | 9.80E+09 | 5.70E+09 | 1.79E+10 | 6.30E+09 | 3.95E+09 |
| **Treated Subject** | **D0** | **D5** | **D12** | **D19** | **D33** | **D97** |
| Ciprofloxacin (16 μg/mL) | 1.25E+09 | 2.33E+09 | 4.30E+08 | 8.80E+08 | 9.35E+08 | 5.35E+08 |
| Tetracycline (32 μg/mL) | 6.35E+08 | 7.05E+08 | 9.00E+06 | 3.50E+08 | 3.85E+08 | 6.15E+08 |
| Erythromycin (32 μg/mL) | 3.45E+08 | 7.40E+08 | 3.49E+08 | 4.95E+08 | 2.95E+08 | 3.25E+08 |
| Metronidazole (16 μg/mL) | 1.79E+08 | 8.80E+08 | 3.51E+08 | 1.96E+08 | 6.10E+08 | 1.92E+09 |
| Vancomycin (32 μg/mL) | 1.23E+09 | 2.91E+09 | 1.30E+07 | 1.63E+09 | 1.75E+09 | 1.49E+09 |
| Chloramphenicol (32 μg/mL) | 0.00E+00 | 2.30E+05 | 2.30E+03 | 0.00E+00 | 0.00E+00 | 0.00E+00 |
| Clindamycin (16 μg/mL) | 5.10E+07 | 2.95E+08 | 3.78E+08 | 1.15E+08 | 4.65E+07 | 8.45E+07 |
| Cephalexin (32 μg/mL) | 6.05E+09 | 2.81E+10 | 1.25E+09 | 9.75E+09 | 1.12E+10 | 1.70E+10 |
| Cefuroxime (32 μg/mL) | 6.50E+09 | 2.59E+10 | 1.35E+09 | 1.02E+10 | 6.25E+09 | 1.36E+10 |
| Cefotaxime (16 μg/mL) | 9.85E+08 | 1.25E+10 | 9.50E+08 | 9.50E+09 | 4.35E+09 | 8.90E+09 |
| Cefepime (16 μg/mL) | 4.62E+09 | 2.71E+10 | 2.60E+09 | 1.63E+10 | 1.15E+10 | 1.68E+10 |
| Dicloxacillin (32 μg/mL) | 7.35E+09 | 1.02E+10 | 2.20E+09 | 8.65E+09 | 7.95E+09 | 1.48E+10 |
| Ampicillin (32 μg/mL) | 1.24E+10 | 2.38E+10 | 1.65E+09 | 1.13E+10 | 1.22E+10 | 1.19E+10 |
| Gentamycin (32 μg/mL) | 1.54E+10 | 2.65E+10 | 2.65E+09 | 1.90E+10 | 1.11E+10 | 1.73E+10 |
| Sulfamethoxazole (64 μg/mL) | 3.65E+09 | 2.15E+10 | 9.50E+08 | 1.60E+10 | 4.15E+09 | 3.95E+09 |
| Pipercillin (32 μg/mL) | 1.79E+09 | 8.00E+09 | 1.05E+09 | 8.50E+09 | 9.30E+09 | 1.39E+10 |
| No Antibiotic | 1.69E+10 | 2.80E+10 | 3.80E+09 | 2.11E+10 | 1.78E+10 | 2.07E+10 |
