## Supplementary figures and images for "Antibiotic Treatment Drives the Diversification of the Human Gut Resistome"

### Abundance variation for beta-lactamases.

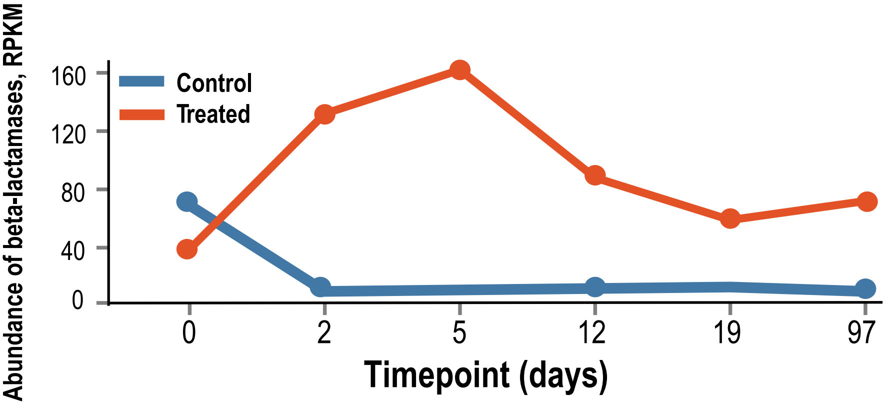

### Abundance variation for various types of antibiotic resistant genes in A. beta-lactam plates and B. non-beta-lactam plates.

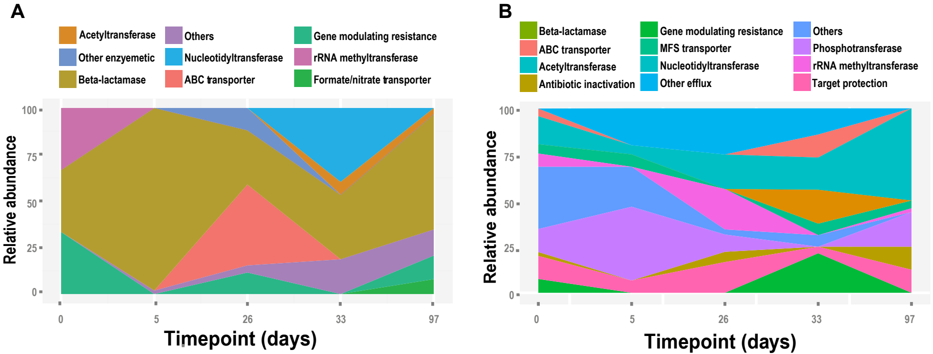

### Correlation between the variation of HGT potentials and HGT or SNP rate.

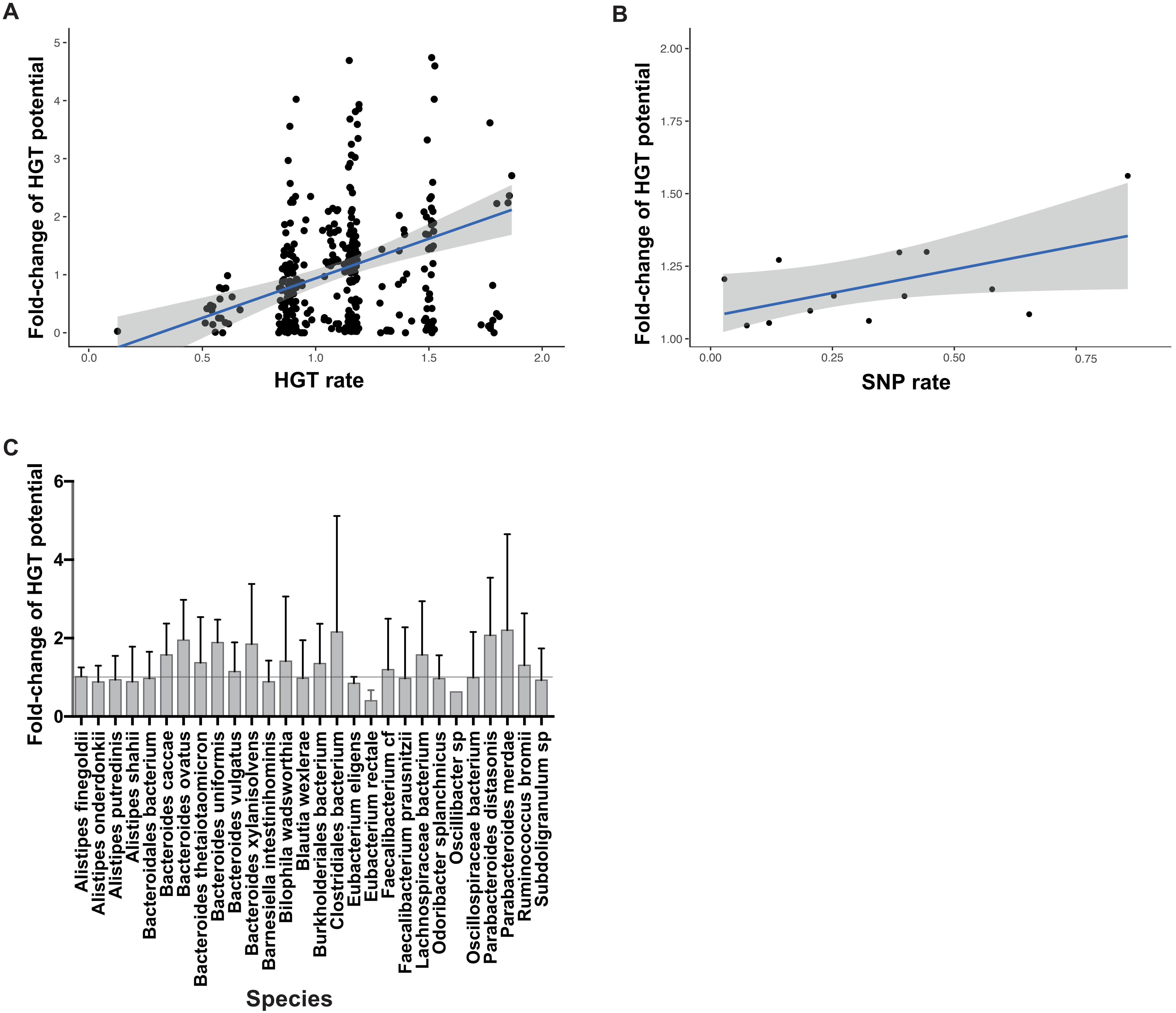

### Temporal variation of the nucleotide diversity in the predicted ARGs in both treated and control subject.

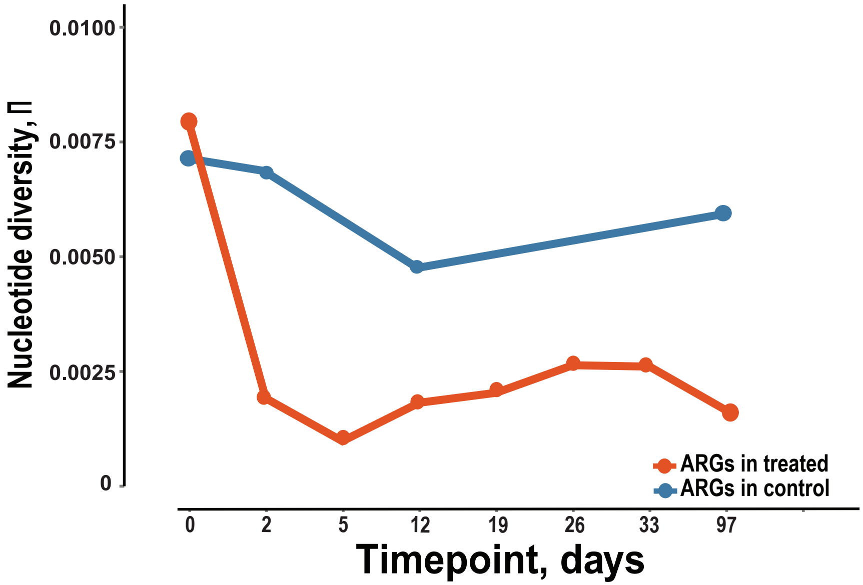

### The average Bray-Curtis distance based on the ARGs abundance profile between individuals.

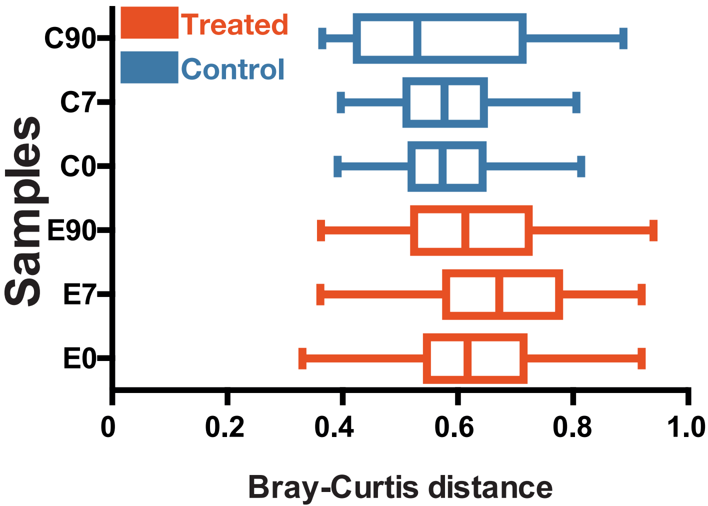

### The average Morisita-Horn dissimilarity of the resistant and controlled communities cultured.

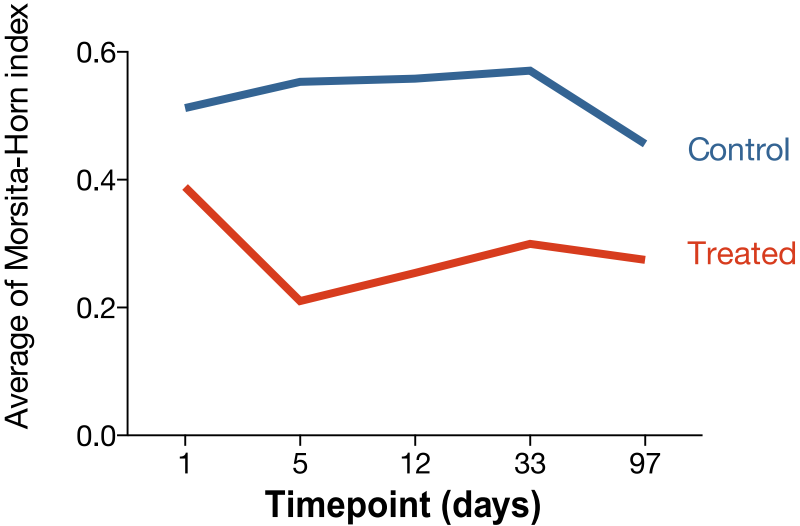

### The pairwise Morisita-Horn dissimilarity (beta-diversity) between cultured communities with antibiotics from all time points.

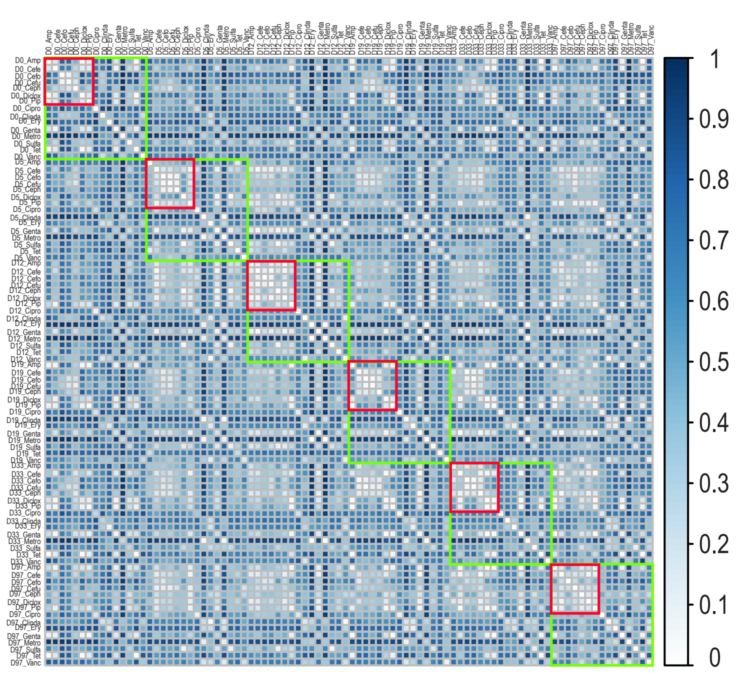

### The percentage of genes with recent HGT signatures based on public database.

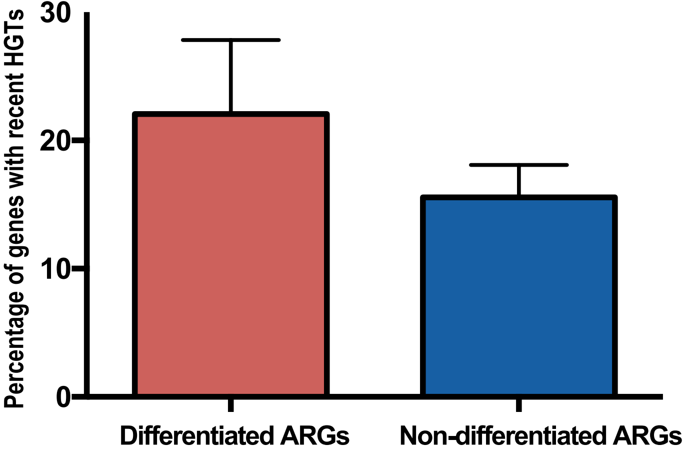

### The phylogenomic tree of A. dominant strain in species Ruminococcus bromii and B. all dominant genotypes of antibiotic resistance genes.

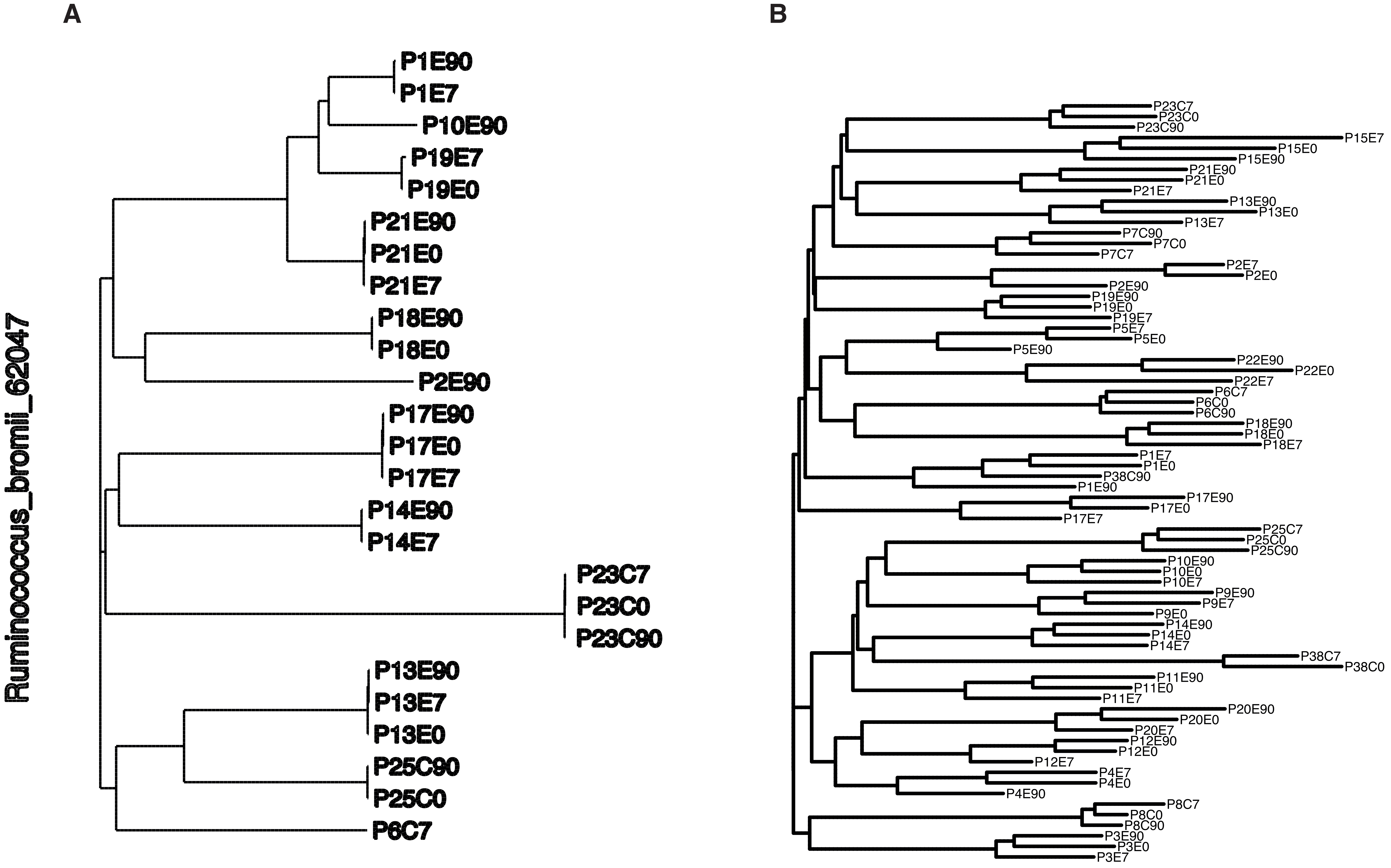

### Variation of the genome-wide diversity over time in the control subject.

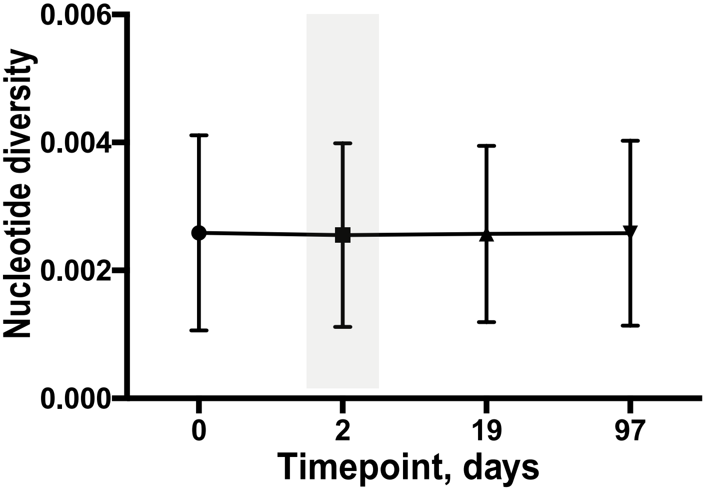

### Variation of the HGT potential over time for two types of ARGs in the control subjects.

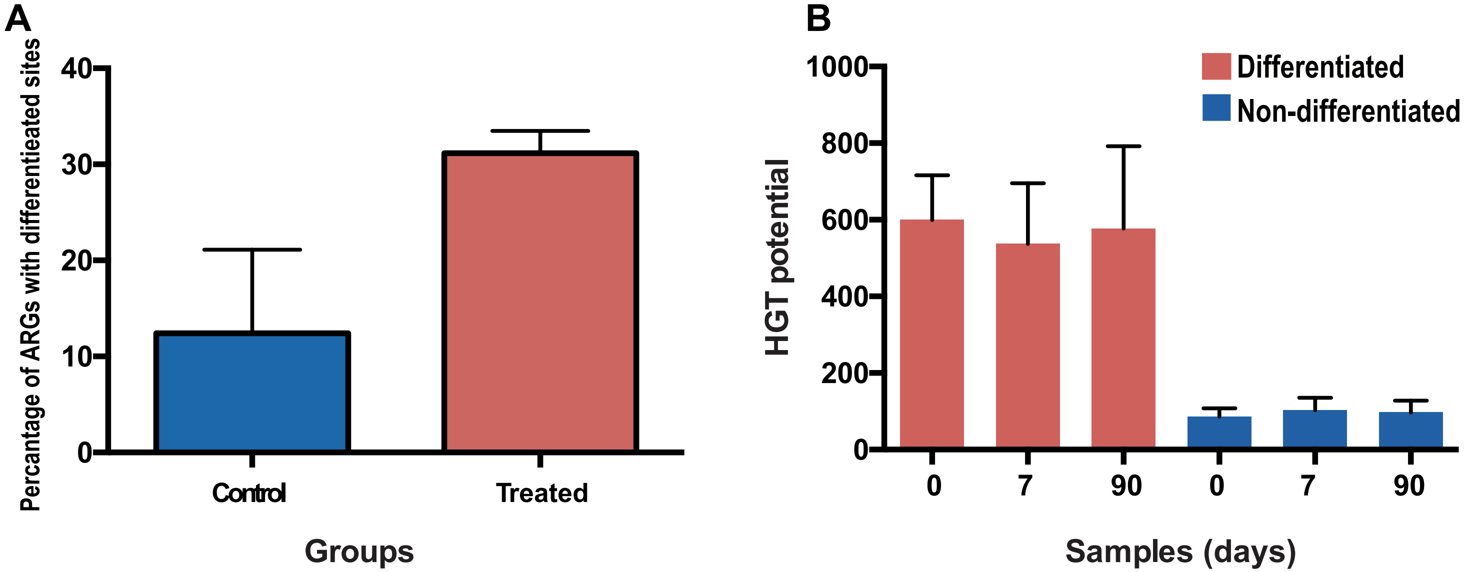

### Variation of the relative abundance and HGT potential of the functionally selected ARGs.

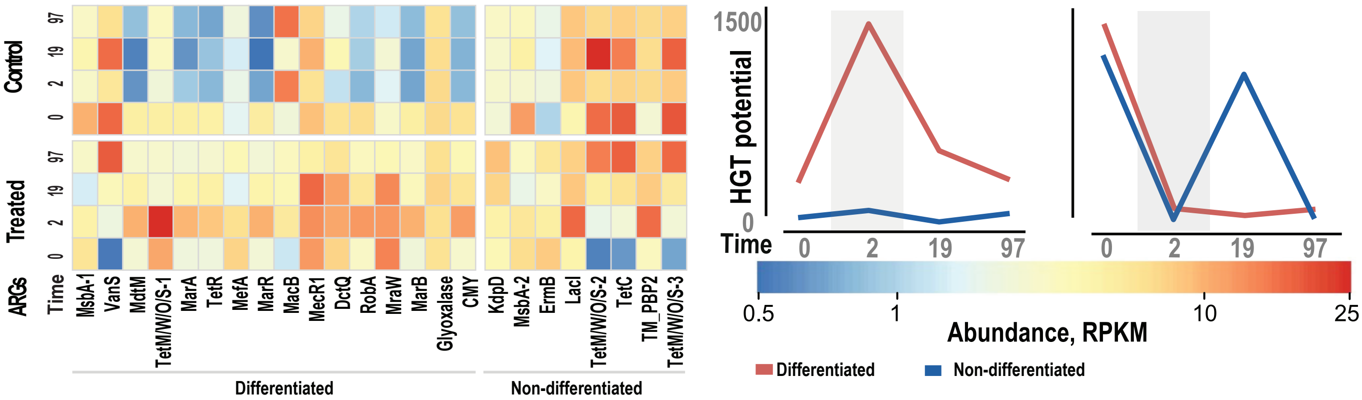
